# Exploring fractal dimensions on ultrasound of gonadal images for sex determination in shortnose sturgeon (*Acipenser brevirostrum*)

**DOI:** 10.1101/2024.10.31.621316

**Authors:** Keyvan Balazadeh, Matthew K. Litvak

**Affiliations:** Department of Biology, Mount Allison University New Brunswick, Canada

**Keywords:** shortnose sturgeon, ultrasonography, ultrasound, fractal dimensions, texture analysis, gonads

## Abstract

Sturgeon are often viewed as monomorphic species because they often lack external features that allow differentiation between sexes. Current practices for sexing sturgeon rely heavily on surgical invasive procedures to visually examine gonadal tissues. An alternative which is widely used in different species of Acipenserids is the use of ultrasound to sex them. The challenge with ultrasound is that it requires an experienced operator to successfully sex the fish by recognizing relevant patterns in the structures of gonads. The objective of this study is to attempt to lay the groundwork for potential systematization of sexing techniques using a portable ultrasound as a medium. We used texture analysis software based on lacunarity measurements and fractal dimensions to determine whether male gonads, female granular tissue (GT) and female pinheads were significantly different from each other using two different resolutions on the ultrasound machine. Male gonads and GT were significantly different from pinheads in lacunarity measurements (p=0.001) using 10-12Mhz general resolution imaging. Fractal dimensions did not result in any significant differences. Lacunarity has the potential to determine sex based on gray level-co-occurrence matrices and the software is freely available. We have also developed a relative probability table based on the data gathered on lacunarity in this study available as supplementary material for quick reference.

## 1. Introduction

Acipenseriformes is a family of fish, comprised of 27 different species of sturgeon and paddlefish, most of which are considered at risk or endangered (IUCN 2016). Their caviar is highly valued, which has generated much commercial interest and aquaculture development since the 1980’s (Pikitch *et al*. 2005; Raymakers 2006; Engler and Knapp 2008; Wuertz *et al*. 2009; Litvak 2010; Kynard *et al*. 2016).The goal of most sturgeon aquaculture activities is to grow females for caviar production and remove males from the production stream. Seventeen species of sturgeon along with two Polydontidae species and their hybrids are currently under commercial production and/or development throughout the world (Bronzi and Rosenthal 2014). Shortnose sturgeon has generated interest for aquaculture development in North America because of its small size and early maturation compared to most sturgeons.

Shortnose sturgeon are endemic to the East coast of North America, ranging from the St. Johns River in Florida, to the Saint John River in New Brunswick (Vladykov and Greeley 1963). They were listed as endangered in the US in 1967 (Department of the Interior *et al*. 1973; NOAA 1988) and as species of special concern in Canada in 1980 (COSEWIC 2005, 2015). There are no commercial fisheries for shortnose sturgeon but they are caught as by catch in the American shad and Atlantic sturgeon fisheries (Collins and Smith 1993; Kynard 1997; Collins *et al*. 2000; Bahn *et al*. 2012). Despite their status as endangered, populations continue to decline (COSEWIC 2015). This makes shortnose sturgeon of special interest for aquaculture development for both commerce and conservation. Given that caviar prices are constantly rising (Pikitch *et al*. 2005; Engler and Knapp 2008), aquaculture ventures, when managed properly, can be highly profitable. This will help to satisfy demand for caviar in the world market. The major obstacle in managing sturgeon aquaculture is the ability to sort cultured stock by sex, and doing so at the earliest possible time is key to maximizing profits (Feist *et al*. 2004). Space is one of the biggest constraints in sturgeon aquaculture, so having the ability to sort and remove males from the stock is highly desirable and increases profitability.

Shortnose sturgeon, like other Acipenserids, were thought to be sexually monomorphic (Dadswell 1979; Hurvitz *et al*. 2007; Keyvanshokooh and Gharaei 2010; Kynard *et al*. 2016), which implies that there are no observable morphological differences between sexes. However, we have found that at the adult stage external differences in head morphology do exist and that they are sexually dimorphic. This is a step forward, but we still need to determine gonad stage in a culture setting. Gonadal differentiation may start at approximately 7 months, but spermatogonia and oogonia may not be present until 36 months (Flynn and Benfey 2007). Females may spawn every 3-5 years after reaching maturity (Dadswell 1979) at approximately 18 years of age. In culture, this has been reduced to 6-12 years of age.

Current practices for sex determination in sturgeon rely mostly on surgical procedures; e.g., borescopes, gonadal biopsy, endoscopes and/or laparoscopy (Moghim et al. 2002; Colombo et al. 2004; Hernandez-Divers et al. 2004; Wildhaber et al. 2005; Bryan et al. 2007; Hurvitz et al. 2007; Chebanov and Galich 2009; Divers et al. 2009; Keyvanshokooh and Gharaei 2010; Masoudifard et al. 2011; Petochi et al. 2011; Matsche et al. 2011; Novelo and Tiersch 2012; Munhofen et al. 2014; Rzepkowska et al. 2014; Falahatkar and Imanpour 2014; Webb et al. 2019). Ultrasound is also commonly used; success rates range from 86-97% depending on the user’s experience (Table 1). Unfortunately, developing user competency for sexing sturgeon through ultrasonography takes time. Species of sturgeon on which ultrasound techniques have been applied include shovelnose sturgeon (*Scaphirhynchus platorynchus)*, white sturgeon (*Acipenser transmontanus*), Siberian sturgeon (*A. baerii*), Russian sturgeon (*A. gueldenstadtii*), Adriatic sturgeon (*A. naccarii*), Beluga sturgeon (*Huso huso*), Lake sturgeon (*A. fulvescens*), pallid sturgeon (*Scaphirhynchus albus*), and stellate sturgeon (*A. stellatus*). This method has yet to be applied to shortnose sturgeon in an objective manner.

**Table 1.** A review of methods and success rates in literature using ultrasonography on sturgeon. In methods U = Ultrasonography, E = Endoscopy, B = Blood parameters and S = sex steroids. Probe type L means linear transducer.

| Species of Sturgeon | Author(s) | Year | Methods | Probe Type | Sex Accuracy | Ultrasonographer skill |
| --- | --- | --- | --- | --- | --- | --- |
| Stellate | Moghim *et al.* | 2002 | U | L | 97.2% Overall | Experienced users identified gonads |
| Shovelnose | Colombo *et al.* | 2004 | U | L | 86% Overall | Experienced ultrasonographer identified gonads |
| Pallid | Wildhaber *et al.* | 2005 | U, E | L | Stage dependent | Skill not specified |
| Shovelnose | Wildhaber *et al.* | 2006 | U, E, B | L | Stage dependent | Skill not specified |
| Shovelnose | Bryan *et al.* | 2007 | U | Unspecified | N/A | Descriptive analysis of egg size |
| Russian and Stellate | Chebanov and Galich | 2009 | U | L | N/A | Descriptive user analysis - Experienced |
| Adriatic and Siberian | Petochi *et al.* | 2011 | U, E, S | L | Stage dependent | Skill not specified |
| Beluga | Masoudifard *et al.* | 2011 | U | L | 97% Overall | Skill not specified |
| Beluga Juvenile | Esmailnia *et al.* | 2019 | U, S hormones | L | 81% | Skill not specified |
| Chinese | (Du *et al.* 2017). | 2017 | U, hormones, S | Curved | 67-97% Stage dependent | Skill not specified |
| Various | Novelo and Tiersch | 2011 | U | N/A | Varied | Descriptive review of papers |
| White—wild and hatchery | Maskill *et al.* (Maskill *et al.* 2022) | 2021 | U, E, S, histology | L | 57-70% | Skill not specified—done by committee of 3 |
| Persian | Vajhi *et al.* | 2013 | U | L | N/A | Descriptive analysis of ovaries and testes – Skilled user |
| Siberian | Munhofen *et al.* | 2014 | U | L | 88% Overall | Skill not specified |
| Lake | Chiotti *et al.* | 2016 | U | L | 88-96% | Skill not specified |

The goal of our study is to develop an objective approach to ultrasound image interpretation to predict sex of sturgeon. Grey scale mapping of pixels from ultrasound images allows us to generate measures of surface roughness translating into texture for the gonads. We developed values for texture of gonadal images using fractals and lacunarity (Mandelbrot 1983, 1994). Fractal dimension analysis has been used in eels, and cod (McEvoy et al. 2009; Müller et al. 2016) and is widely used in medicine (Iversen and Nicolaysen 1995; Glenny *et al*. 2000; Karch *et al*. 2003; Marxen and Henkelman 2003; Al-Kadi and Watson 2008; Cheng *et al*. 2010; Valentin *et al*. 2011; Jurczyszyn and Osiecka 2012). Here we compare lacunarity and fractal values between male and female tissue types so that ideally a user with no prior experience with ultrasound image interpretation can determine sex of a shortnose sturgeon.

## 2. Methods

Aquaculture fish were sampled at Breviro Caviar’s hatcheries in Charlo and Pennfield, New Brunswick. Another group of fish from the DFO Mactaquac hatchery, held for our our lab, were sampled, as well as a group of wild caught fish from the Saint John River near Hartt Island (45°58’08.2” N 66°44’38.5” W), New Brunswick. Female sex and reproductive stage were confirmed through either terminal sampling or biopsies with core sample removals for visual inspection. All males were wild caught during spawning season; sex was confirmed by the presence of milt. Females sampled at Breviro Caviar were all a minimum of 6 years of age and weighed 3-10kg. Males were of unknown age, but had an average weight of 3kg.

Images were collected using a portable ultrasound machine (SonoSite, M-Turbo, FujiFilm) equipped with a rectangular transducer capable of scanning from 6-15MHz (model: HL-50). The fish were sampled using general resolution (GEN) which ranges from 10-12MHz along with an additional sample taken using the highest resolution (RES) at 12-15Mhz. Depth of penetration is affected by frequency; lower frequencies penetrate deeper, but with lower resolution.

### 2.1. Ultrasound sampling and classification

The fish were ultrasound scanned by placing the probe between lateral scutes 2-3, 3-4, 5-6, 6-7, and 7-8 (Fig 1). The sections between the scutes were sampled ventrally and then laterally on both the left and right sides of the fish. Frontal views were taken transversely between each scute section. Ultrasound images were taken at a maximum depth setting of 4.0-4.9cm. Brightness and contrast settings were adjusted using the analog dials and varied depending on backlighting required for the user to see the images properly (normally at a bright setting at a high contrast). Only images that lacked aberrations were used. Aberrations result from shadowing caused by penetration problems between scutes, improper probe placement, lost pixels, and spotted smoothing of images while the probe is in movement. The most difficult samples to acquire are generally from females with granular tissue (developing oocytes, earliest female stage, normally pink sandy texture) and pinheads (small oocytes generally >0.5mm to 1mm and white in color). It is very hard to obtain females in these specific reproductive stages, since they typically have mixed tissue that contains small pinheads with oocytes under 1mm diameter. To simplify female classification, if no oocytes were present in any form, they were considered to have granular tissue as long as the coloration and texture matched the reference image (Fig 2); pink coloration with a highly-textured surface. Measurement of the granular tissue oocytes was not possible due to their lack of structural integrity. When samples were removed most of them would dissociate rapidly. Any oocytes greater than 1mm diameter were considered stage 1 or more developed oocytes. Males were classified by their white coloured testes and smoothness.

**Figure 1.**
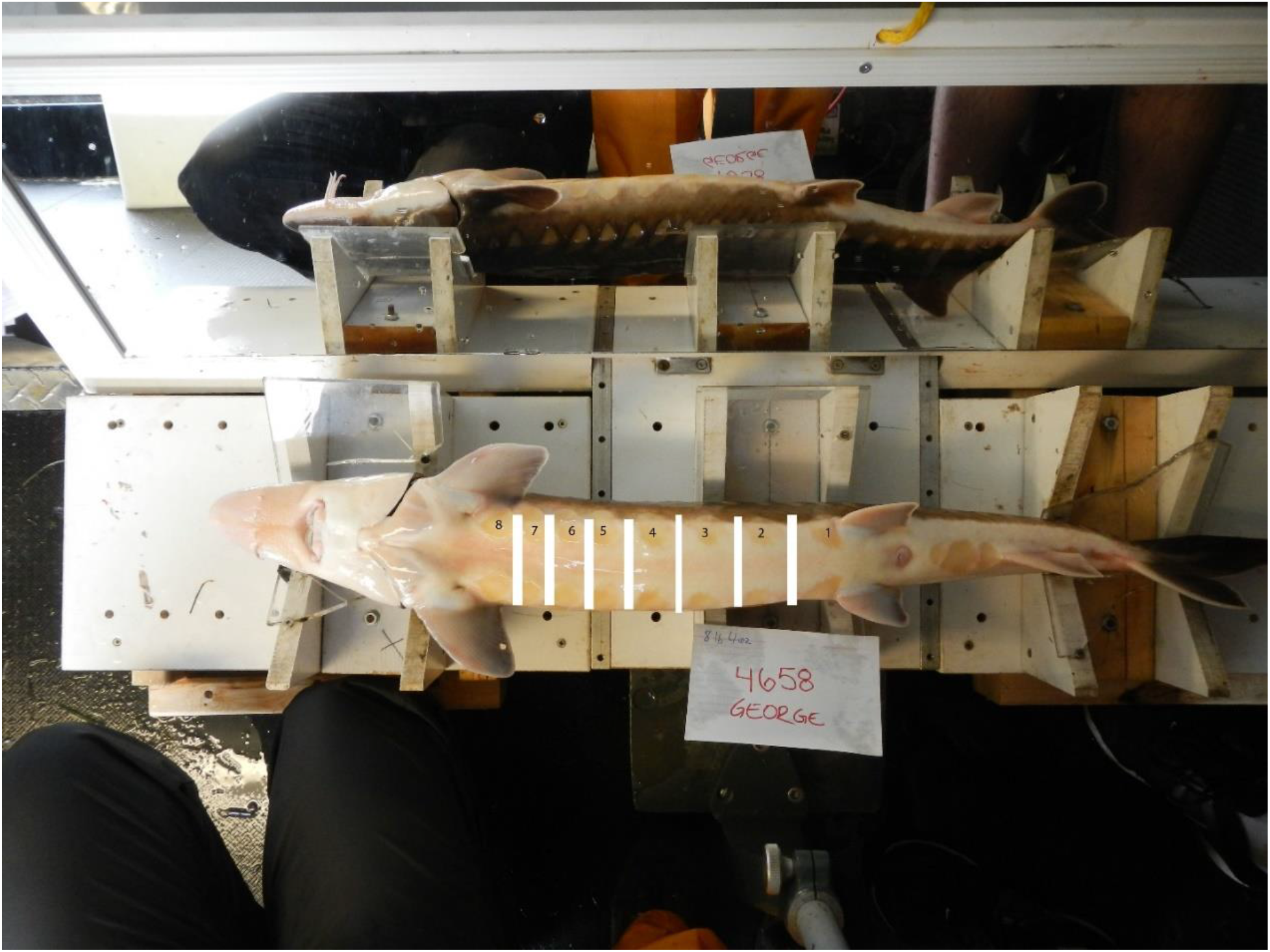
Ventral view of shortnose sturgeon. The white markings represent where the probe was to be placed for imaging. Plexiglass miniature V-troughs were adjust for the fish’s size. For reference, the scutes are numbered 1-8 posterior to anterior.

**Figure 2.**
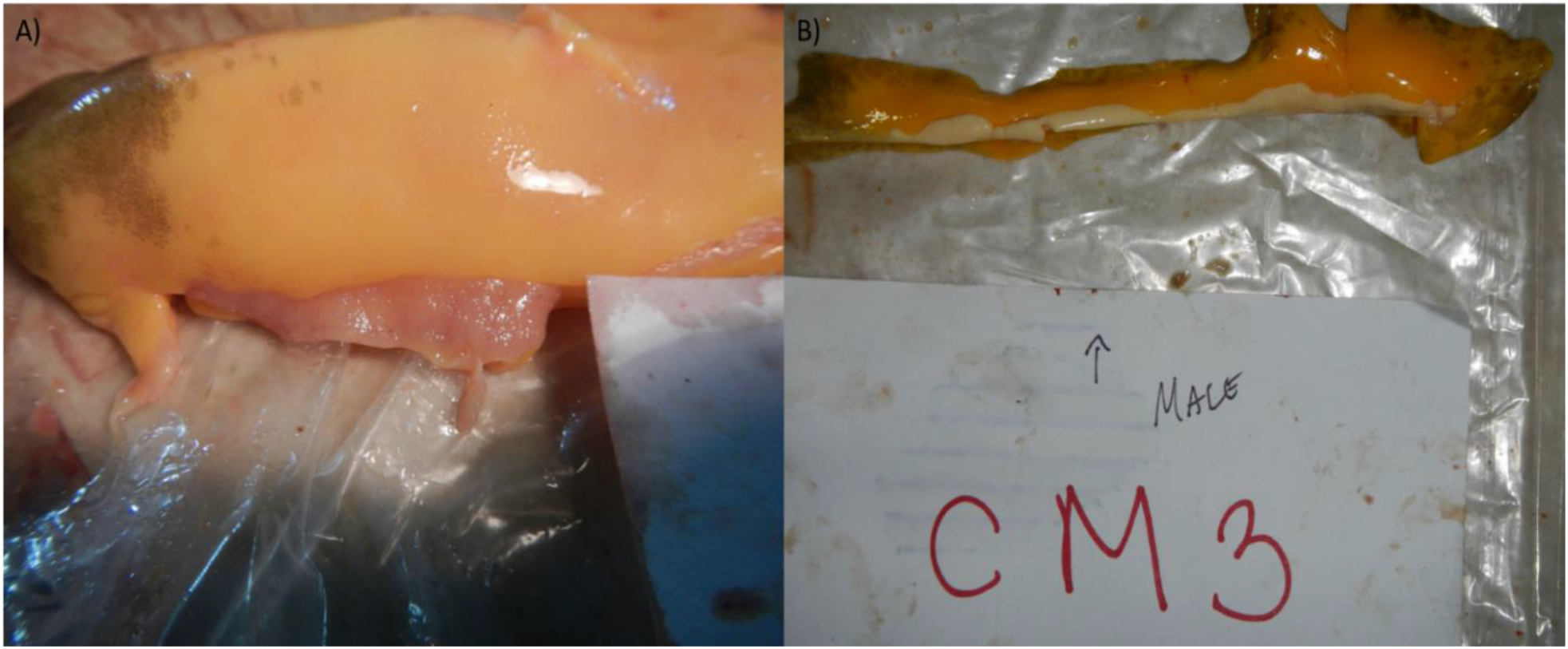
Differentiating between male and female gonad tissues from shortnose sturgeon. A) Female granular tissue, identified by the pink tissue. Yellow tissue with black coloration is fat. B) Male gonad is identified by the white tissue. Yellow tissue is fat

Fish were tagged using a numbered “spaghetti” Floy™ tag. This allowed for identification of individuals sampled at the aquaculture facility to have their sex confirmed during harvesting. The entire ultrasound and biopsy sampling process resulted in fish being exposed to air for no longer than 5 minutes. No fish were lost because of our sampling procedures.

### 2.2 Image Analysis

Ultrasound images, in BMP format, were organized corresponding to individual fish tags. This enabled individuals to be tracked throughout the sex confirmation process. Images showing gonads were collected and analyzed per scute for each fish. Images were then individually labeled with a fish ID and sorted. Regions of interest (ROI) of the images were analyzed for surface texture (Fig 3) for gonad types male, pinheads and granular tissue. Images with stage 1 or more developed oocytes (caviar ready oocytes) were not included in this study because developed oocytes, commonly referred to as eggs, are obvious with ultrasound and can be easily measured and staged by a non-experienced user without requiring any extra analyses or software. We used ImageJ (Rasband 1997) with the plugin “FracLac” (Karperien 1999) to generate fractal values for surface roughness. Surface roughness is visualized based on grey values and their intensities on an image which can be translated into texture (Fig 4). We used the polygon selection tool to determine the ROI of the gonad. This was done so that areas with a high amount of fat were excluded from biasing the fractal measurements. Due to the high amount of sound wave reflections, or echogenicity, the border lines around the gonads were also avoided to prevent skewing measurements. Echogenicity is the ability of a sound or an echo to reflect off a surface. A highly echogenic structure has high reflectivity for sound waves whereas low echogenicity absorbs most of these waves. Images that are hyperechogenic, such as the gonads, are lighter in color whereas images that are hypoechogenic, such as fat, result in darker colors.

**Figure 3.**
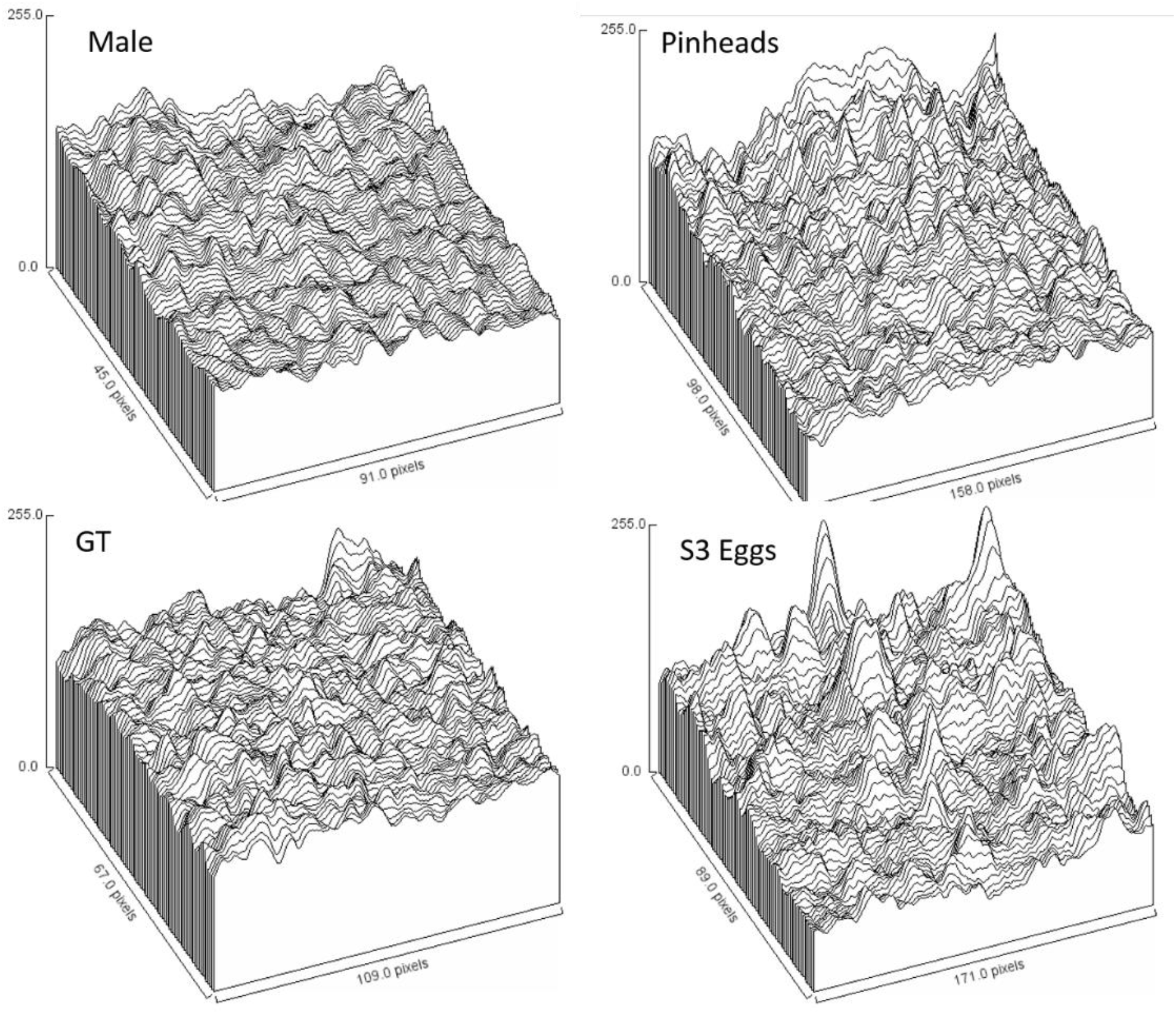
Surface plots generated in ImageJ visualizing surface roughness for various types of gonads. Peaks represent the highest gray values and the valleys are darker pixel values. S3 gonads with eggs were added to see how the peaks become accentuated from pinheads to eggs >2.2mm.

**Figure 4.**
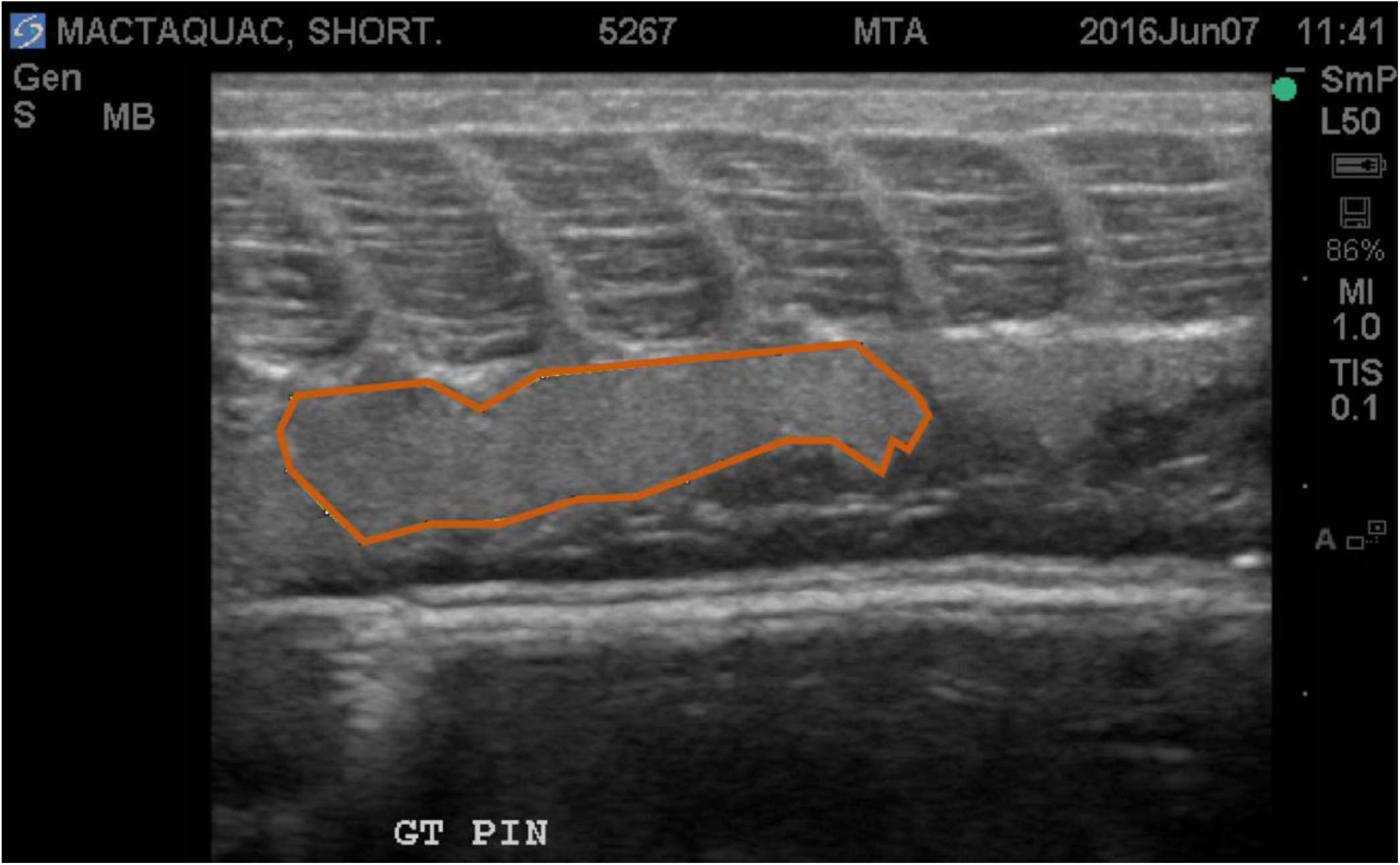
Ultrasound image depicting the sampling area using the polygon tool in ImageJ prior to running FracLac. The total image visible has 307,200 pixels, the enclosed polygon has about 111,290 pixels which is our area of interest.

Once the area was bound, the image was converted into an 8-bit image in ImageJ, cropped to the selected size, and submitted to FracLac. In FracLac, block series were selected with 50 positions which provides 50 iterations of the process. Minimum pixel size was set to 3, which provides a minimum 3×3 pixel box for scanning with 45% of the image minimally sampled. This was the preferred approach because images are 640×480 pixels, which gives a maximum pixel count of 307200 pixels. We used a minimum of 45% of the sampled image results in 138240 pixels available for sampling. This is important because it allows us to understand how much of the image is used to analyze the gonad. By further using 3×3 pixels as a minimum sample size of a box, it ensured that the algorithm did not miss possible pinhead eggs on the female gonads since this size is greater than a full pinhead oocyte. When the image is cropped, and focused on a particular spot on the gonad, the number of total pixels used is relative to that specific area enclosed by the polygon (Fig 3).

Gray 1 differential was selected for image type in FracLac. Color code, lacunarity (a measure of heterogeneity) and draw grids were also selected for graphics options. Once the settings were fixed, the images were scanned in FracLac to extract the values. Values for lacunarity and fractal dimensions were then obtained from the “Box Count Data Per Scan” file. Lacunarity values were then derived using the formulas (Equation 1) described in Karperien (1999) and Al-Kadi and Watson (2008). M and N represent the length and width, in pixels, of the processed image. Λ is lacunarity, ε is box count or scale, g is grid orientation. The basic equation for fractal dimensions (Equation 2) uses “r” as the scaling factor setup in the software while N_r_ represents the number of self-similar shapes or the number of boxes enclosing the image (Al-Kadi and Watson 2008).

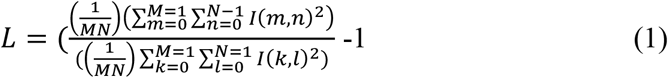

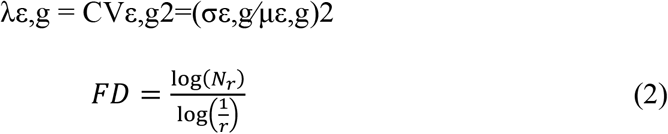

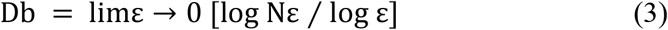

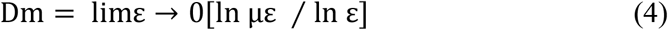

The fractal dimension for box counting, Db, is calculated using the size of boxes and number of boxes calculated in a grid (Equation 3) using the slope of the regression line as the limit. Mass, Dm (Karperien 1999), is calculated by counting the number of pixels from within a box on a grid at a given scale (Equation 4); the limit of ε → 0 is calculated from the regression line of µε and ε. µε is the mean pixels per box at some ε where ε is the box size or scale. These values are used because variance will not be altered with different gray intensity values that may be found with different machines or outputs. This is only true if the sampled image area does not contain parts in which pixels are completely blacked out when they should not be. It is important to select only parts that are actual gonads and not fat or body cavity. Lacunarity uses the gray pixel counts and values and calculates a coefficient of variation. If excessive black pixels that do not correspond to gonads are introduced the lacunarity value becomes exceedingly skewed.

Tests were run to assess the variability of lacunarity using different sized boxes within 10 samples of a sample tile with a homogeneous textured layout. There was little difference in variability (0.001) from a box 3 times than its original size. The area of the upper gonad visible in the screen was also calculated. The entirety of the gonad was measured with a surrounding polygon to calculate visible surface area. The area of the ROI and the area of the total gonad visible was also recorded (Fig 5). This gives an indication of how much of the gonad was measured to obtain the fractal and lacunarity values presented in this study.

**Figure 5.**
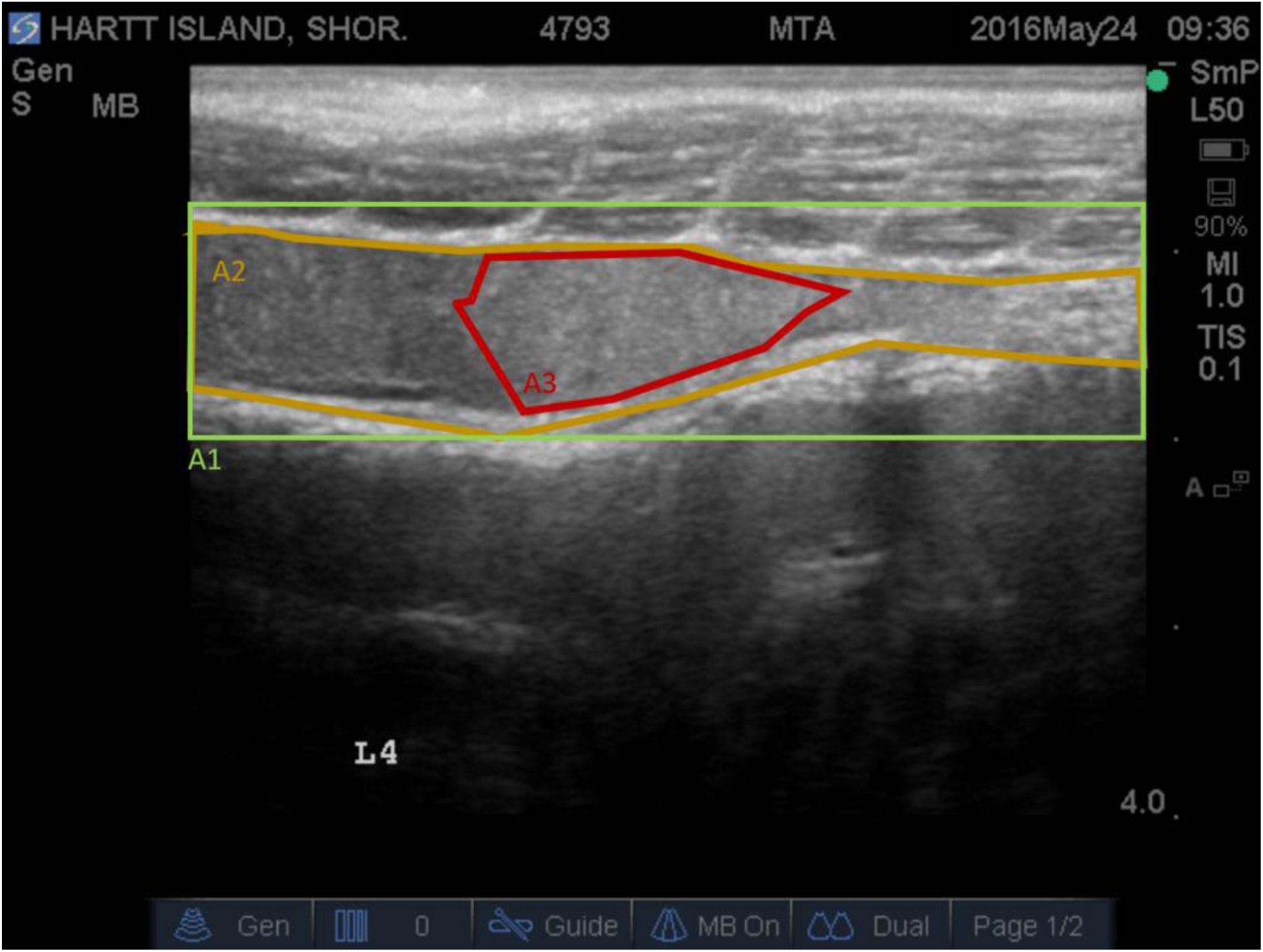
Measured areas of gonadal images. A1) Region of interest containing the visible gonad with a rectangular area. This extends normally from the edge of the myosepta, separating it from the alimentary tract, and extending to the membrane of the lower gonad. This encompasses the entire gonad. Since the gonads are not rectangular, the surrounding rectangular area is a best estimate containing both the upper border with the muscles and the lower border of the gonad with the digestive tract. A2) Entire gonad area, including fat content and aberrations. A3) Bounding polygon representing the area of interest sampled. This contains the lowest number of aberrations and ideally no fat

### 2.3. Statistical Methods

One-way ANOVAs were run on both resolutions for lacunarity, Db and Dm. Since data contained mixed individuals for both resolutions, we could not statistically compare between them. Since variables/fractals were generated on the same image, a correction for experiment-wise error rate was calculated by using a sequential Bonferroni correction of alpha for the ANOVAs (Holm 1979). In this case the ANOVA with the lowest p-value is compared to alpha/3 (0.0167), the second lowest p-value to alpha/2 (0.025) and the lowest p-value to alpha=0.05. Tukey’s HSD was run to test for significant differences when ANOVAs were significant. Assumptions of normality and homogeneity of variance were assessed with Shapiro Wilk’s test and Levene’s test, respectively. Analyses were conducted using R (v3.3.3) with an RStudio (v1.0.136) interface. A series of power analyses were conducted using G.Power (v3.1.9.2) at 80% power with effect sizes calculated with standard deviations and group means for each variable. This allowed us to estimate sample sizes required for future studies. Three effect sizes were used: a small effect size (0.1), a medium effect size (0.25) and a large effect size (0.4) based on Cohen’s D (Cohen 1992). Additionally, a set of sample estimates were added for effect sizes of 0.1, 0.25 and 0.4 at 80% power using three groups at a 95% confidence interval for ANOVA.

## 3. Results

A total of 58 fish (25 males and 33 females) were sampled using the general view (10-12Mhz) or GEN. Of the females, 23 were granular tissue stage and 10 were pinheads. A total of 39 fish were sampled using the high-resolution view (12-15Mhz) or RES. Of those, there were 20 males, 11 females in the granular tissue stage and 8 females containing pinheads. All lacunarity and fractal data were normal based on Shapiro Wilk’s tests (p-values for LacGEN= 0.843, LacRES = 0.474, DmGEN = 0.234) except for Db on GEN and RES (p = 0.038, p = 0.0002 respectively) along with Dm on RES (p = 0.011), which were skewed because of two samples. A visual assessment of normality was conducted which checked out and the two skewed samples were kept in. All lacunarity and fractal data passed Levene’s test for homogeneity of variance (the lowest p-value was 0.59 for LacRES). No transformations were conducted on the data as ANOVAs are robust to violations of parametric assumptions (Zar 2010).

Lacunarity in the general resolution (GEN) was the only value with a significant difference between gonad types [F(2,55) = 7.716, p = 0.0001]. As a reference, values for oocytes larger than pinheads were included as a subsample, these are Stage 1+ eggs which are greater than 1mm in diameter. The average fractal values for stage 1 along with more developed egg stages resulted in Db = 1.372, L= 0.1288 and Dm = 0.572 in the GEN resolution images. Since females containing eggs are easily identifiable by nearly anyone using an ultrasound without any experience, these values are to be used as reference only since they are beyond the scope of this study. As reference, a set of images was added corresponding to lacunarity values and their respective gonad types including stage 3 eggs (Fig 6).

**Figure 6.**
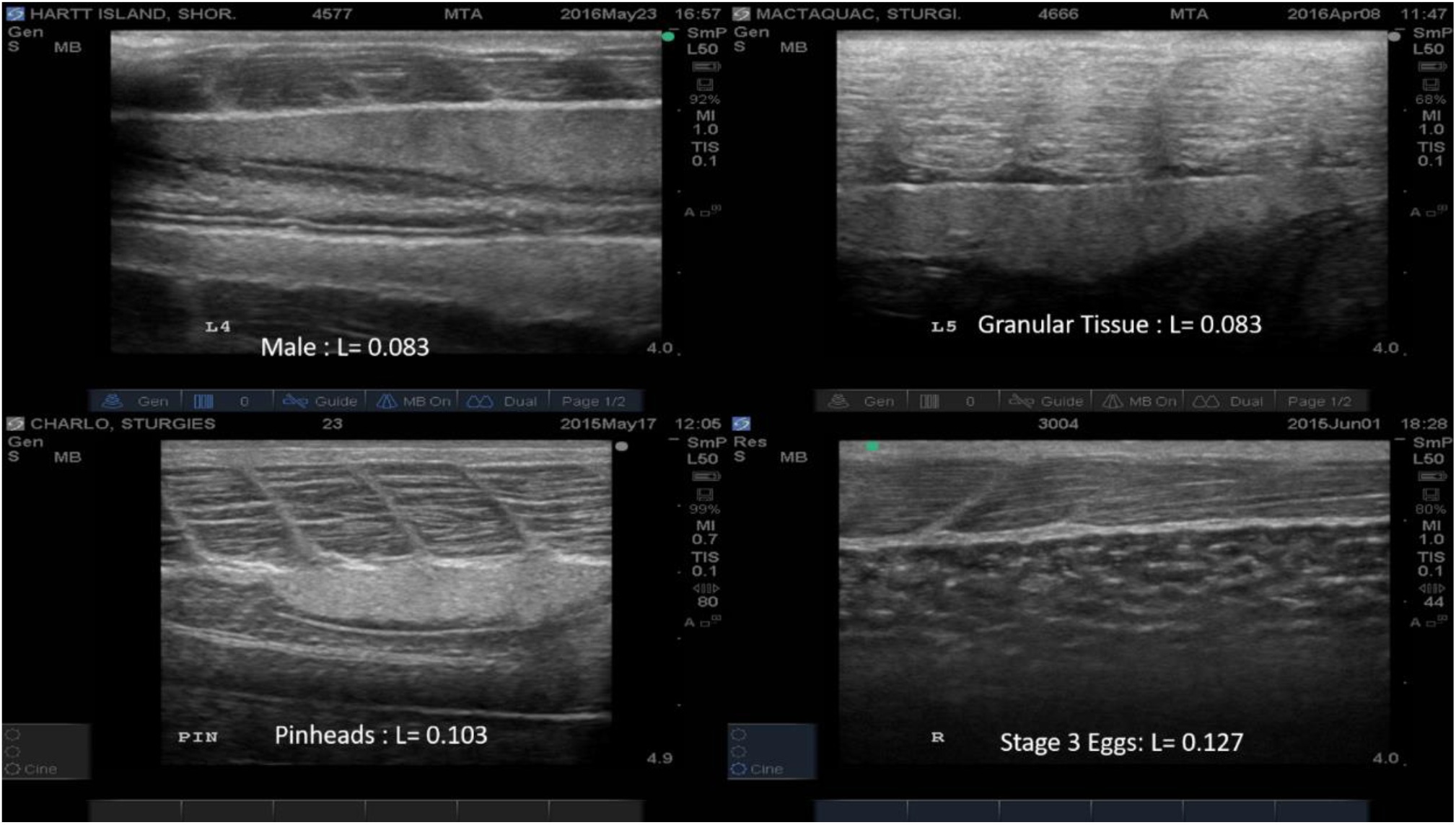
Sample of gonad types with corresponding lacunarity values on ultrasound images

The comparison of lacunarity values of all three gonads types was significant for only GEN resolution images. No significant differences were found on the high-resolution test RES images although it is nearly significant [F(2,36) =3.004, p = 0.0622]. This may be a result of a lack of power due to low sample size of fish using RES resolution. Visualizing eggs is easier in the GEN resolution than the RES resolution setting. The reason is because oocytes tend to become hidden due to insufficient penetration of sound waves when fat content is high, it is also very common for multiple sizes of oocytes to be present. The GEN resolution has better penetration and most of these issues do not apply.

The major difference in lacunarity is visible between pinheads and granular tissue (Tukey’s test, p=0.003), and pinheads and male gonads (p=0.002). Differences in means are notable for both GEN and RES images in lacunarity, however, the GEN resolution has a less variable and a larger difference when compared to RES resolution images (Fig 7).

**Figure 7.**
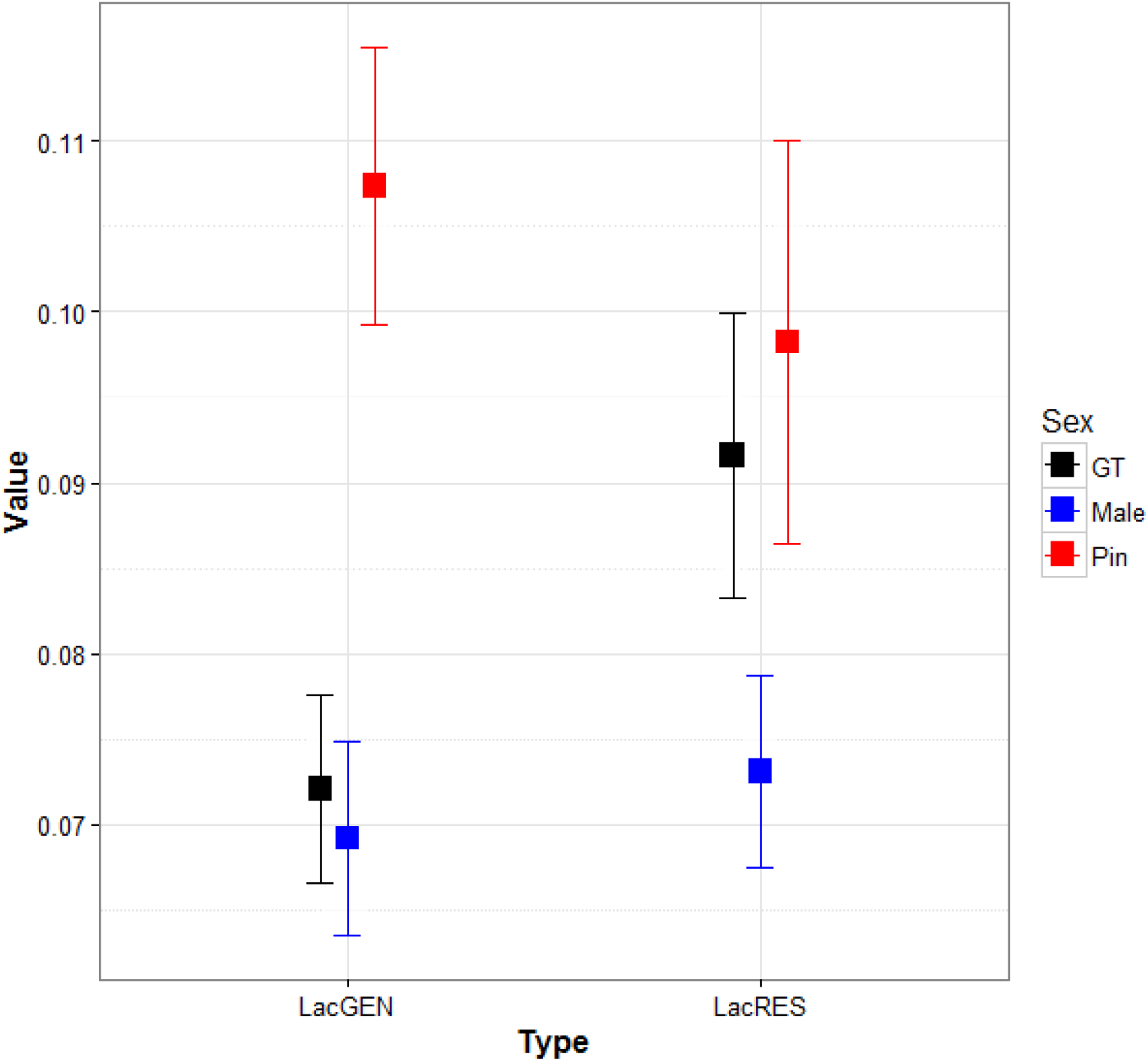
Results of mean differences in lacunarity between each gonad type for both types of resolution. Value is a unitless measure of lacunarity

Fractals DbGEN [F (2,55) = 1.043, p = 0.359], DbRES [F (2,36) = 0.522, p = 0.598], DmGEN [F (2,55) = 0.092, p = 0.912] and DmRES [F (2,36) = 0.268, p = 0.766] were tested in the same manner as lacunarity but no significant differences were observed between the gonads at both resolutions.

The level of variation is higher in RES images when compared to GEN images, although the differences are minor (Table 2; Fig 7). Even though they are not comparable statistically, pinheads and granular tissue distribution are less separated and more normalized in the GEN resolution than in RES. This may indicate that although some oocytes may be hidden in higher resolutions, resulting in a less accurate depiction of pinhead values, the males are detected better. Males also showed a high degree of variability present in the RES images suggesting possible variation of stages within the male gonads.

**Table 2.** Summary of coefficients of variation of lacunarity values for each gonad type.

| **Gonad Type** | **Mean CV GEN** | **Mean CV RES** | **SD CV GEN** | **SD CV RES** |
| --- | --- | --- | --- | --- |
| **GT** | 0.103 | 0.144 | 0.084 | 0.214 |
| **PIN** | 0.055 | 0.058 | 0.027 | 0.031 |
| **MALE** | 0.112 | 0.082 | 0.104 | 0.061 |

Overall, it is easier to establish regions of interest in females than males due to the amount of discernable gonad in ultrasound images (Table 3; Fig 6). Compared to females, males have higher variation present when it comes to image brightness, that is, males tend to have a more variable range of gonad echogenicity which is seen by a larger range in brightness. There is certainly user bias included in this, but when compared to males, female gonads tend to have a larger area of visible gonad without aberrations making it easier to select the ROI.

**Table 3.** Summary of gonad area calculated from the total area of the gonad including fat content ^*^(A2) and the percentage of that gonad area used in the region of interest ^*^(A3).

|  | **GT ROI area (mm<sup>2</sup>)** | **PIN ROI Size (mm<sup>2</sup>)** | **Male ROI Size (mm<sup>2</sup>)** |
| --- | --- | --- | --- |
| **Average** | 17.48 | 19.49 | 16.04 |
| **SD** | 3.00 | 7.25 | 6.56 |
| **Mean % used from total gonad** | 78.14 | 80.57 | 58.86 |
| **SD % used from total gonad** | 11.09 | 17.47 | 20.59 |

\*(A2) Refer to Fig 5. Represents the entire gonad area, including fat content and aberrations.
\*(A3) Refer to Fig 5. Bounding polygon representing the area of interest to be sampled. Contains lowest number of aberrations and ideally no fat.

The power analysis suggested that over 200 samples would be required to detect a significant difference between gonad types in fractals on both resolutions (Table 4). The one exception is LacRES which would only require approximately 72 samples to detect a difference between gonad groups at a nearly large effect size at 0.4 (calculated effect size = 0.378).

**Table 4.** Power analysis based on F-values obtained from one-way ANOVAS. Confidence intervals for all samples was 95%. The calculations were done using means for each gonad type along with their respective standard deviation, these are 3 groups. Power was set to 80% to estimate sample size. Effect size varied for each different calculation based on standard deviation.

|  | **Samples required** | **Calculated effect size from SD** | **Critical F** |
| --- | --- | --- | --- |
| **LacRES** | 72 | 0.378 | 3.1 |
| **DbGEN** | 234 | 0.205 | 3.03 |
| **DbRES** | 324 | 0.173 | 3.02 |
| **DmGEN** | 354 | 0.166 | 3.02 |
| **DmRES** | 675 | 0.119 | 3 |

In the supplementary material section, we have added two tables (Table S1; Table S2) that may be useful as a guide to sex fish with given lacunarities. There are probabilities attached to the tables in regards of how likely it is to have a male or granular tissue female at a given stage and likewise for a female to be a pinhead or more developed female using a t-distribution generated from our data.

## 4. Discussion

Ultrasonography has been used to determine sex for a number of sturgeon species (Table 1). However, to our knowledge all analyses to date have been subjective; they have depended on user interpretation of ultrasonic images. While this approach has been successful, it depends on user capability and experience. There has been a great deal of research on the development of objective approaches to image interpretation in medicine (e.g.: Iversen and Nicolaysen 1995; Glenny *et al*. 2000; Karch *et al*. 2003; Marxen and Henkelman 2003; Al-Kadi and Watson 2008; Cheng *et al*. 2010; Valentin *et al*. 2011; Jurczyszyn and Osiecka 2012). We sought to determine the potential of image analytic techniques for interpretation and classification of sex and stage in shortnose sturgeon. This approach removes the effect of observer experience and potentially provides a consistent and repeatable approach to interpretation of sex and reproductive stage. Fortunately, there have been tremendous advances in image analysis over the past 20 years and much of the software is available freely. Here we used fractals (lacunarity within the FracLac plugin in ImageJ) to examine image surface roughness to successfully and objectively discriminate between male and female tissue. This approach frees-up the requirement of an experienced ultrasound operator to view images to sex and stage sturgeon in aquaculture.

When comparing different gonads, females with stage 1+ oocytes have very distinct lacunarity values, which makes them easy to define. Pinhead lacunarity values are also distinct from granular tissue and male gonads. However, the problematic lacunarity values are from male tissue and female granular tissue. Vajhi *et al*. (2013) claimed that early reproductive stages can be differentiated between male and female gonads, their approach used an experienced ultrasonographer and did not use an objective computer assisted approach. Unfortunately, lacunarity alone will not allow us to determine sex at early stages of female gonadal development prior to the pinhead stage, leaving granular tissue and male tissue as the hardest stages to discern.

There is a large amount of variation within granular tissue as well as male tissue in lacunarity values. Some of this variation in lacunarity signals can be attributed to user error while selecting the regions of interest for analysis. Another possibility is potential variability in male reproductive readiness and development (Chebanov and Galich 2009; Golpour *et al*. 2017) affecting these values. There are also reported differences of female oocyte development in white sturgeon in which oocytes may remain in a dormant state, lacking yolk, for periods of a year or more (Doroshov *et al*. 1997). This may have been the reason we saw variability in lacunarity values we observed within the granular tissue stage. This variability has also been seen in other studies sexing sturgeon (Moghim *et al*. 2002; Wildhaber *et al*. 2007; Chebanov and Galich 2009; Masoudifard *et al*. 2011; Petochi *et al*. 2011; Munhofen *et al*. 2014; Chiotti *et al*. 2016).

Furthermore, the use of ultrasound allows for quick identification of developed eggs in sturgeon, however care must be taken to use the appropriate frequencies in order to avoid losing information. The limitation of high-resolution imaging (12-15MHz range) is depth of penetration in visualizing gonads. We found the penetration was not enough to record pinheads and, in some cases, small stage 1 eggs. This was particularly evident with recently developing eggs in gonads after confirming samples by internal inspection. Some of these samples would contain a few eggs encrusted within layers of fat and developing granular tissue which can be hard to visualize with ultrasound. Obtaining samples at specific stages on females is a very complex and difficult task unless thousands of fish are collected. These limitations in identifying female stages correctly are expected because while higher frequencies lead to higher resolution, it reduces penetration (Masoudifard *et al*. 2011).

Fractals have been used in an attempt to identify sex in cod by examining gonads (McEvoy *et al*. 2009), resulting in an accuracy of 80% in sex determination. However, in our study, we did not find fractals Db or Dm to have any detectable significant differences between male and female gonads. We think this is because fractals require shapes to be present to find patterns and create an accurate fractal dimensional value. Lacunarity examines the gaps between gray values on a surface; if there are grey value peaks there are also grey value valleys, which create these gaps. In other words, it examines the smoothness or the heterogeneity of the surface that is imaged, not specific patterns. A different approach to examining image surface texture is by analyzing gray level co-occurrence matrices which are appropriate in our images and can be done using GLCM (Cabrera 2003) in ImageJ. The difference between GLCM which has been used in aquatic species such as eels (Müller et al. 2016) and lacunarity is that GLCM is susceptible to differences in brightness and contrast settings since it cannot correct for any changes in brightness and contrast. Lacunarity is a more complex way of determining gray values since it is based on the variation for these grey values and not just the intensity of the peak values. We ran preliminary tests to see if there were differences in lacunarity and GLCM between 1-6 points of contrast in ImageJ. Lacunarity was barely affected (0.0001 change) whereas GLCM values were greatly affected primarily in the “contrast and inertia” variables (7 points of difference) and a change of 15 points in the “variance” variable. GLCM is susceptible to changes in sampling conditions while lacunarity is not. We sampled at different times in the day, weather conditions, location and penetration settings. That is, both in the wild and in varying hatchery conditions requiring the brightness settings of the machine to be constantly adjusted. Furthermore, the Sonosite M-Turbo uses a set of analog dials to adjust brightness and contrast settings making it very hard to track and quantify brightness and contrast settings. Selecting an ultrasound machine with digital versus analog contrast and brightness setting adjustments is important, since this limitation made following the procedure of eliminating background colour thresholding employed by McEvoy *et al*. (2009) impractical.

The use of fractal dimensions and lacunarity in oncology relies on high resolution images from CT (Al-Kadi and Watson 2008) or clinical ultrasound machines (Cheng *et al*. 2010; Valentin *et al*. 2011). Fractal dimensions are highly affected by image resolution (McEvoy *et al*. 2009), so images generated from higher resolution CTs are much more amenable to fractal analysis. In our study, the maximum resolution obtained using the SonoSite M-Turbo was 640×480 pixels, whether it is at 6mhz or 15mhz. This is a limitation of midrange portable ultrasound models. Given that our results depicted differences between pinhead stages compared to granular tissue and male gonads in lacunarity, perhaps such high resolutions are not necessarily needed. This does not remove the possibility that the results could be enhanced with higher pixel resolution in the near future or with more powerful ultrasound machines.

Based on our methods lacunarity remained the best predictor for sex on the three gonadal stages we studied. The power analysis conducted on fractals Db and Dm showed that the required sample sizes to find significant differences unrealistic. Since image quality varies between ultrasound machines and the combination of probes used, lacunarity should be further examined using a combination of different devices.

The supplementary tables provided (Table S1;Table S2) can be used as a rough guide of what to expect for lacunarity values and the probability of being a male or female. If fish, in a facility, were sorted every 6 months using these parameters it may help select females with very little effort as they develop from granular tissue stage to pinheads. Since females spawn every 3-5 years (Dadswell 1979), using a two-step or a four-step sorting trial within 2-3 years, most females would be selected from the group. Our results suggest that lacunarity holds promise for objective classification of gonad types without the reliance upon user experience. This also suggests that there is potential for development of automated sexing systems, such as AI deep learning approaches, for sturgeon with ultrasonography.

## Acknowledgements

We would like to thank NBIF/NSERC/CRD for grants to MKL and Breviro Caviar for their support for this study. This study would have had significantly fewer samples if Breviro Caviar did not allow the use of their facilities and fish to examine. We would also like to thank all the MTA Litvak Lab research assistants who helped during field collection. Thanks to Christine Gilroy for help she provided both as assistant and lab technician in the field and in the lab. Special thanks to the staff at the Mactaquac DFO hatchery for being always helpful.

## Supplementary Tables

**Table S 1.** Probability of a gonad type being classified as a pinhead or more developed eggs based on a t-distribution generated from lacunarity values.

| Lacunarity | Probability of<br>being Pinhead % |
| --- | --- |
| 0.01 | 0.21 |
| 0.02 | 0.38 |
| 0.03 | 0.72 |
| 0.04 | 1.36 |
| 0.05 | 2.58 |
| 0.06 | 4.85 |
| 0.07 | 8.90 |
| 0.08 | 15.62 |
| 0.09 | 25.71 |
| 0.1 | 38.99 |
| 0.11 | 53.98 |
| 0.12 | 68.34 |
| 0.13 | 80.04 |
| 0.14 | 88.30 |
| 0.15 | 93.50 |

**Table S 2.** Probability of being classified as either a male and/or female with granular tissue based on a t-distribution at given lacunarity values. The lower the lacunarity, the higher the chance of being a male. Note these values group male and GT together as one since there was no detectable difference between them in this study.

| **Lacunarity** | **Probability %** |
| --- | --- |
| 0.01 | 98.48 |
| 0.02 | 96.58 |
| 0.03 | 92.93 |
| 0.04 | 86.73 |
| 0.05 | 77.39 |
| 0.06 | 65.06 |
| 0.07 | 50.81 |
| 0.08 | 36.45 |
| 0.09 | 23.84 |
| 0.1 | 14.15 |
| 0.11 | 7.62 |
| 0.12 | 3.73 |
| 0.13 | 1.67 |
| 0.14 | 0.68 |
| 0.15 | 0.26 |

